# Facing small and biased data dilemma in drug discovery with federated learning

**DOI:** 10.1101/2020.03.19.998898

**Authors:** Zhaoping Xiong, Ziqiang Cheng, Chi Xu, Xinyuan Lin, Xiaohong Liu, Dingyan Wang, Xiaomin Luo, Yong Zhang, Nan Qiao, Mingyue Zheng, Hualiang Jiang

**Affiliations:** Shanghai Institute for Advanced Immunochemical Studies, and School of Life Science and Technology, ShanghaiTech University, Shanghai 200031, China; Drug Discovery and Design Center, State Key Laboratory of Drug Research, Shanghai Institute of Materia Medica, Chinese Academy of Sciences, 555 Zuchongzhi Road, Shanghai, 201203, China; University of Chinese Academy of Sciences, No.19A Yuquan Road, Beijing 100049, China; School of Information Science and Technology, University of Science and Technology of China Hefei, 230000,China; Laboratory of Health Intelligence, Huawei Technologies Co., Ltd, Shenzhen, 518100, China

**Author notes:** Corresponding Author. (Hualiang Jiang). (Mingyue Zheng). Authors contribute equally (Zhaoping Xiong, Ziqiang Cheng, Chi Xu).

## Abstract

Artificial intelligence (AI) models usually require large amounts of high-quality training data, which is in striking contrast to the situation of small and biased data faced by current drug discovery pipelines. The concept of federated learning has been proposed to utilize distributed data from different sources without leaking sensitive information of these data. This emerging decentralized machine learning paradigm is expected to dramatically improve the success of AI-powered drug discovery. We here simulate the federated learning process with 7 aqueous solubility datasets from different sources, among which there are overlapping molecules with high or low biases in the recorded values. Beyond the benefit of gaining more data, we also demonstrate federated training has a regularization effect making it superior than centralized training on the pooled datasets with high biases. Further, two more cases are studied to test the usability of federated learning in drug discovery. Our work demonstrates the application of federated learning in predicting drug related properties, but also highlights its promising role in addressing the small data and biased data dilemma in drug discovery.

## Introduction

Current artificial intelligence (AI) has high requirements for data both in terms of quality and quantity to achieve good predictive performance. Data acquisition difficulties and data biases in the measurement of scientific tests have significantly limited AI’s power in drug discovery^1–3^. Data acquisition challenges come from time-consuming and expensive processes of data-generation and consequent confidentiality, especially in the later stages drug development such as data about drug pharmacokinetics, safety, and efficacy profiles. Taking ADME/T (Absorption, Distribution, Metabolism, Excretion and Toxicity) properties as an example, such data are usually highly standardized and of good quality, and would contribute to better predictive model and generate larger added value when used for modeling. However, few of these properties are exposed to the latest deep learning models due to confidentiality^4,5^, which can be considered as an enormous loss for drug development. Not only the data acquisition difficulties resulting from confidentiality, the data biases in the measurement of scientific tests also perplex AI for drug discovery^6^. It is common to see, for example, a specific molecular property has large discrepancy in recorded values from different sources, even under the same measurement of the same scientific tests. The discrepancy in recorded values is usually considered to come from data biases, because a recorded value in each data source is obtained from repeating measurements and the effect of variance has been reduced to minimum. In conventional machine learning paradigm, the discrepancy in different sources will usually be uniformed by taking the mean, median or a majority vote, which might bring the value closer to the “ground truth”. However, for practitioners who generated those data may only care about the recorded value in their own experimental setting-ups for reference (i.e. whether the structure modification will lead to the optimization of a property), rather than the absolute “true” value. Therefore, a shared global model for all data sources might not form good instructions to practitioners. Federated learning emerges as a new machine learning paradigm provide viable solutions for data acquisition and data bias problems faced by AI drug discovery by keeping confidentiality and customizing model for users.

Federated learning represents a scenario where multiple clients can train a model collectively without sharing raw data^7–10^. The original idea dates back to 2016, in the context of the enactment of GDPR (General Data Protection Regulation) in Europe, users gain more control over the use of personal data, which challenges many companies that rely heavily on selling Ads based on users’ personal data. McMahan et al. from Google proposed federated learning and a year later initially applied it in Gboard, the keyboard on android phones^7,8^. Therein, federated learning was adopted to train, for example, a next word prediction model crosses many phone devices without uploading users’ data to central servers. This process has improved users’ input experience and preserved users’ privacy. Noteworthily, aside from training a single global model collectively on clients’ datasets, it becomes viable for each client to have a customized model in federated learning^9,10^. Therefore, the problem of discordant records in centralized machine learning turn into an intrinsic feature of federated learning^10^. Federated learning has attracted substantial attention and has found more and more applications in a much broader areas^13–17^, which is also a promising approach to satisfy the needs of drug discovery^18^ but yet to be investigated and tested. Drug discovery has similar request to protect the confidential or IP-sensitive data, and at the same time, to extract the maximum information/knowledge present within such data by machine learning. Moreover, given the high biases in drug discovery related data, customizing a model for each client is appealing for personalized prediction as done in Gboard^8^.

Here, we setup a general federated learning framework for drug discovery (Fig. 1) and tested it on FATE (Federated AI Technology Enabler)^19^, an open-source project aiming at providing a secure computing framework for federated learning. Different from previously mentioned Gboard cross-device federated learning application that trained cross millions of phone devices, federated learning for drug discovery is another setting trained cross data silos, which is termed as cross-silo federated learning. In this setting, there are a coordinator server and several collaborators instrumented with federated learning client program. These clients in collaboration can be big pharmas, biotech startups or even academic labs having their own data silos.

**Fig. 1.**
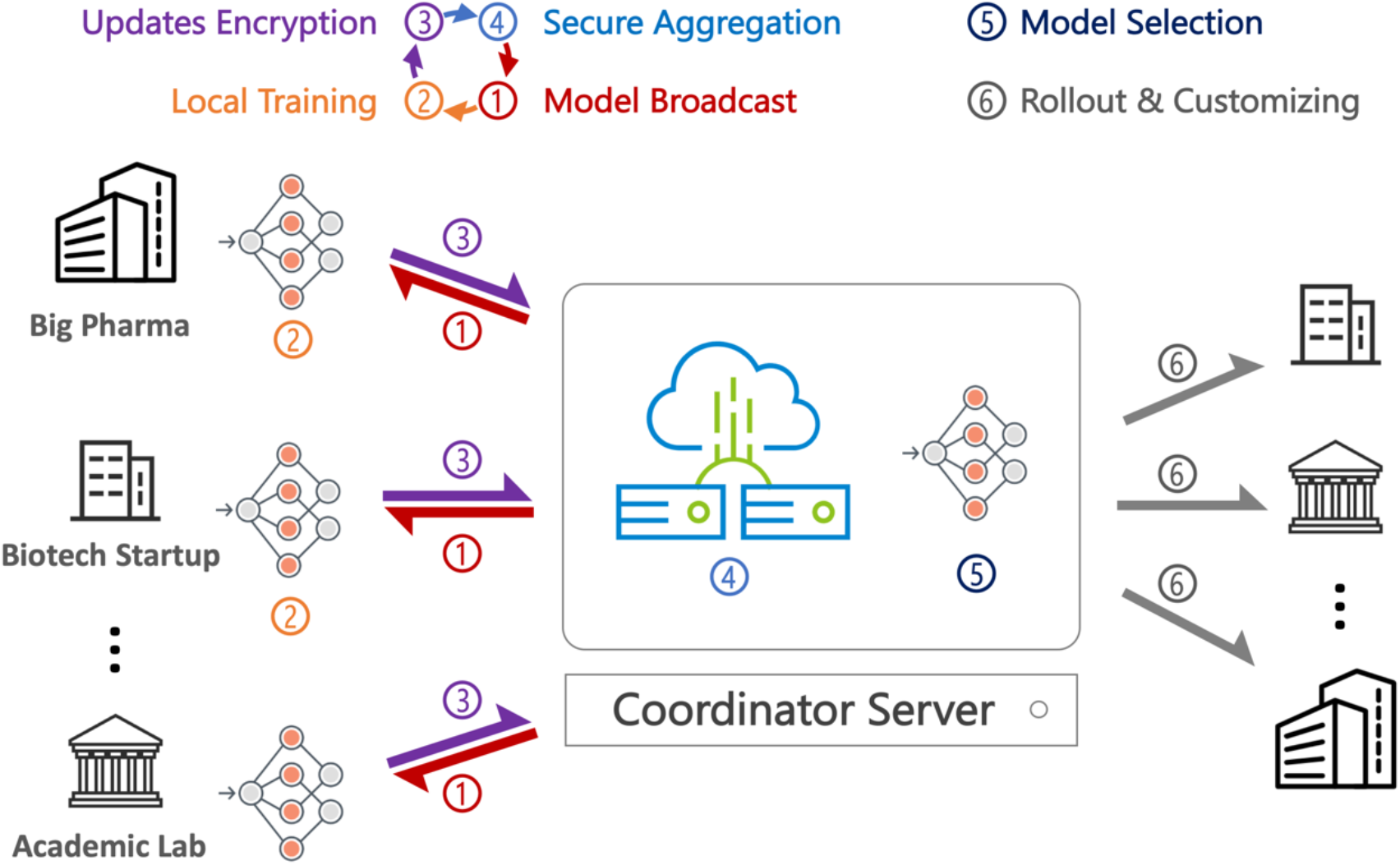
The life cycle of a federated learning system for drug discovery. In federated training: 1) the coordinator server broadcast the latest shared global model to each client; 2) the client locally computes the model updates, 3) encrypts and uploads the model updates; 4) finally, the coordinator server aggregates all the encrypted model updates securely and uses them to update the shared global model for the next round of training. After the training is done, the best model is selected for rollout and might be customized for the users who have their own labeled data.

During each round of cross-silo federated training, 1) the coordinator server broadcast the latest shared global model to each client, 2) each client locally computes the model updates by executing the training program, 3) and encrypts and uploads the model updates under a secure aggregation protocol, 4) finally, the coordinator server aggregates all the encrypted model updates securely and uses them to update the shared global model. Figure 1 illustrates the life cycle of federated learning system for drug discovery. Many rounds of training will be required until the model converges or meets the criteria for stopping, which might be the metrics that does not improve within given rounds on a shared dataset on the coordinator server or a held-out validation dataset on each client. The best model is then selected for rollout. For users who want to use the model for prediction, they can use the selected shared model directly. Alternatively, for users who own plenty of labeled data themselves, they can opt to instrument the federated training program and locally update the model (without uploading updates to coordinator), thus obtaining a customized model. Model customization is a common application and should be very practically useful.

In this work, we simulated cross-silo federated learning processes in three use cases: solubility prediction, kinase inhibitory activity prediction and hERG liability prediction. The datasets in these use cases show variance in the chemical space of compounds covered, measurement methods, experimental conditions, nonstandard representations and size of data. These real-world drug property datasets from different sources represent non-identical data distributions at different clients, from which we would like to investigate how drug discovery projects can benefit from federated learning. Tested with different network structures and federated aggregation algorithms, federated model can always outperform models built on only individual datasets. So we can rely on federated learning to build more predictive model if possible.

## Results

### Facing non-IID data

In conventional centralized machine learning application for drug discovery, to include more data, researchers collect data from different sources, and assume the data are independent identically distributed (IID). The IID sampling of the training data is important to ensure that the stochastic gradient is an unbiased estimate of the full gradient. However, the assumption is usually violated due to the high data biases introduced in the measurement of scientific tests, which are conducted by different people in different experimental setting-ups. As shown in Table 1, we collected water solubility datasets from different sources and some of the shared molecules between datasets can have distinct recorded values. For example, dataset F1 and dataset C2 have 4 shared molecules, the values of those molecules in two datasets have a Mean Absolute Deviation (MAD) of 1.52. The large MAD of shared molecules between datasets may signify the data distribution varies across datasets in some extent. In conventional machine learning, even a model predicts perfectly right on one dataset, it can’t predict well on another due to the violation of IID. Aside from a shared global model for all datasets, it is viable for federated learning to customize a model for each dataset, which is practically useful to deal with biased/Non-IID data.

**Table 1.** Data statistics and the mean absolute deviation of the values for the compounds shared among different sources

|  |  |  |  |  |  |  |  |
| --- | --- | --- | --- | --- | --- | --- | --- |
| F1 | **6110 (0.00)** |  |  |  |  |  |  |
| F2 | 763 (0.27) | **4650 (0.00)** |  |  |  |  |  |
| F3 | 72 (0.35) | 215 (0.12) | **2603 (0.00)** |  |  |  |  |
| F4 | 520 (0.24) | 952 (0.04) | 136 (0.15) | **2115 (0.00)** |  |  |  |
| C1 | 52 (0.19) | 194 (0.06) | 20 (0.22) | 185 (0.06) | **1291 (0.00)** |  |  |
| C2 | 4 (1.52) | 29 (0.09) | 59 (0.11) | 12 (0.12) | 7 (0.22) | **1210 (0.00)** |  |
| C3 | 195 (0.31) | 545 (0.12) | 32 (0.28) | 486 (0.09) | 174 (0.02) | 8 (0.19) | **1144 (0.00)** |
|  | F1 | F2 | F3 | F4 | C1 | C2 | C3 |

\*The numbers outside the parentheses are shared molecules between two datasets, and numbers inside are the mean absolute deviation (MAD) of LogS of these shared molecules.

### Tuning model update frequency

A six-layer DNN is constructed as depicted in Figure 2B, which is a 6-layer multilayer perceptron with neuron numbers of 4096, 1024, 256, 64, 8 and 1 respectively. Compared with conventional centralized model, federated models have the same network architecture except for that after a given epoch of local training, the model updates of each client will be encrypted and uploaded to the coordinator server, and later the coordinator returns a new model to each client. Therein, the model update frequency, i.e. how many epochs should the client run locally before uploading the encrypted model updates, is an influential hyper-parameter. As shown in Table 2, the model updates in every 5 epochs of local training performances yielded the best predictive performance averaged on 5 independent runs.

**Table 2.** Performances of using different federated averaging epochs.

| Client /<br>Test size | Federated averaging epochs |  |  |  |  |
| --- | --- | --- | --- | --- | --- |
|  | Every 1 | Every 5 | Every 10 | Every 15 | Every 20 |
| F1/611 | 0.878±0.022 | **0.864±0.023** | 0.872±0.017 | 0.888±0.013 | 0.870±0.012 |
| F2/465 | 0.543±0.018 | **0.517±0.011** | 0.522±0.007 | 0.519±0.009 | 0.531±0.011 |
| F3/260 | 0.534±0.007 | **0.503±0.011** | 0.511±0.002 | 0.515±0.020 | 0.530±0.006 |
| F4/212 | 0.324±0.037 | **0.296±0.007** | 0.311±0.011 | 0.304±0.009 | 0.310±0.008 |
| C1/258 | 0.299±0.023 | **0.277±0.003** | 0.280±0.004 | 0.283±0.004 | 0.287±0.006 |
| C2/242 | 0.801±0.024 | **0.775±0.004** | 0.789±0.007 | 0.779±0.005 | 0.786±0.005 |
| C3/229 | 0.328±0.021 | **0.313±0.005** | 0.315±0.006 | 0.319±0.010 | 0.322±0.004 |

\*The reported performance is the MAE of LogS on test sets in 5 independent runs. Clients F1-4 are the federated training participants, and have 1/10 of the dataset as the test set (8/10 for federated training and 1/10 for validation), while clients C1-3 participate in customization only, and has 1/5 of the dataset as test set (3/5 for customization training and 1/5 for validation). The best performance on each client is highlighted in bold.

**Fig. 2.**
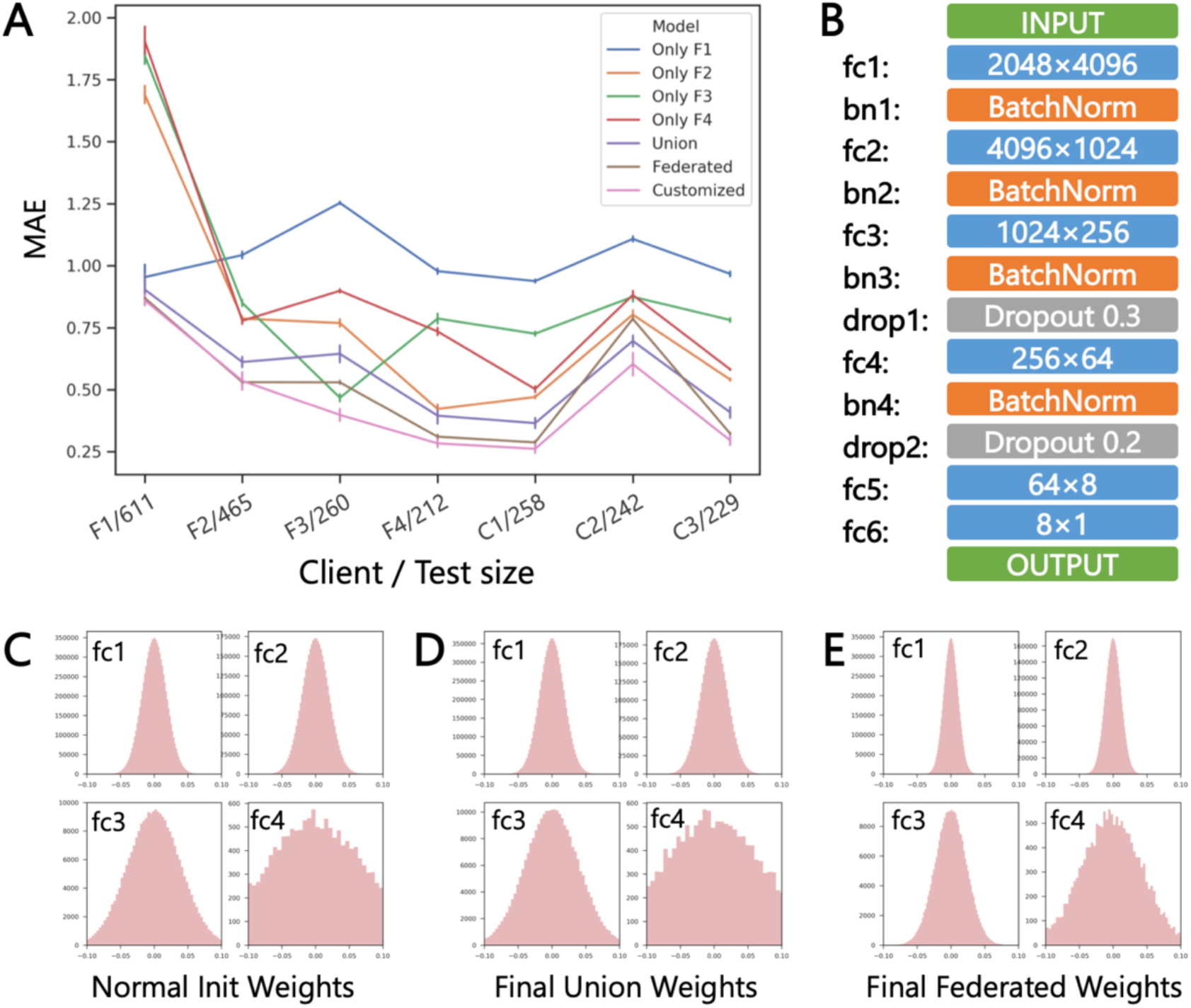
Performance comparisons between federated modeling and centralized modeling. (A) Centralized models are trained on individual clients (Only F1, F2, F3 and F4) or on the union/pooling of datasets F1/F2/F3/F4 (Union), while federated learning models are trained cross clients F1/F2/F3/F4. The MAE of LogS on the test set of each clients are reported. (B) The deep neural network architecture of the model, which is shared by federated modeling and centralized modeling. Fc is fully connected layer with relu activation. (C) When the weights of fc1-4 are initialized with normal distribution, (D) the weight distributions of union model don’t vary tangibly after centralized training, while (E) the weight distributions of federated model vary tangibly with more weights concentrated on 0.

### Performance comparison with centralized baselines

In this study, datasets F1-4 were used for simulating clients who participate in the training process of the federated learning models, and C1-3 for simulating clients who didn’t participate in training but want to customize the federated model with their own data. We compared federated modeling with individualized and centralized modeling baselines (Figure2 and Supplementary Table 1), in terms of the MAE values on the test set of each client. Generally, the sub-models trained on individual datasets achieved higher performance on their own internal test set (i.e., F1/611, F2/465, F3/260 and F4/212), but much lower performance on other test tests, indicating these sub-models can’t generalize well. In contrast, the federated learning model and Union model showed much improved predictive performance on cross client datasets. For clients F1-4, the federated learning model generally yielded lower MAE values than the corresponding sub-models trained locally, and the prediction capability was maintained on tests from external clients C1-3. It is worth noting here that the federated model performed even better than the Union model, in which data from different sources are simply pooled together for training in a non-privacy-preserving way. This is counter-intuitive, to examine their differences in learning, we compared the weight distribution of fully connected layers in the Union model and the shared Federated model (Figure 2C-E). The weight distributions of the Union model basically unchanged after centralized training compared with the initialized weight distribution, while the weight distributions of federated model vary significantly with more weights concentrated on 0. In the same network architecture with the same cohort of parameters, more weights of 0 means that the model is regularized and simpler, which is likely to generalize better^20,21^. This regularization effect explains the side benefit of federated learning when preserving data privacy on training datasets with different systematic biases. As shown in Table 1, dataset F3 showed a larger systematic bias, where the compounds shared with datasets F1 and F4 have averaged MAD values of 0.35 and 0.24, respectively, which may cause the inferior performance of the Union model and the shared federated model when testing on dataset F3. However, when the shared federated model is further fine-tuned with small learning rate locally without uploading the updates, which is a form of customization for the local data, the performance of the federated model can be further improved, especially for datasets with higher biases (i.e., F3 and C2).

### Improved network architecture and aggregation algorithm

To investigate how different network architectures and federated learning aggregation algorithms will influence the performance of federated learning, apart from the previous MLP architecture and FedAvg aggregation algorithm, a residual fully connected neural network (RFCN) architecture^22^ and the FedAMP^23^ aggregation algorithm are also tested. The RFCN model in our experiment is composed of a fully connected layer with 1536 neurons followed by a ResNet of two 2-layer blocks and one single-layer block (Supplementary Figure 2B).

The centralized RFCN model (Union + RFCN) outperforms the federated learning model with MLP and FedAvg algorithm (FedAvg + MLP) on 6 out of 7 clients. This means the centralized MLP model (Union + MLP) tested in the previous section is not a strong baseline and the MLP + FedAvg model outperforms it easily owing to the regularization effect of FedAvg. But with a strong centralized baseline (Union + ResNet), the federated learning usually cannot outperform centralized union model (Table 4 and Table 5).

**Table 3.** Performances of using different network architectures and federated learning algorithm

| Client | Test size | Union | Union | FedAvg | FedAvg | FedAMP | FedAMP |
| --- | --- | --- | --- | --- | --- | --- | --- |
|  |  | + MLP | + RFCN | + MLP | + RFCN | + MLP | + RFCN |
| F1 | 611 | 0.904 | 0.871 | **0.87** | 0.89 | 0.875 | 0.889 |
| F2 | 465 | 0.612 | 0.512 | 0.531 | 0.548 | 0.516 | **0.511** |
| F3 | 260 | 0.645 | 0.505 | 0.53 | 0.55 | 0.495 | **0.482** |
| F4 | 212 | 0.395 | 0.288 | 0.31 | 0.286 | 0.282 | **0.262** |
| C1 | 258 | 0.365 | **0.26** | 0.287 | 0.27 | 0.288 | 0.292 |
| C2 | 242 | 0.697 | 0.75 | 0.786 | 0.808 | 0.795 | **0.775** |
| C3 | 229 | 0.408 | **0.304** | 0.322 | 0.31 | 0.324 | 0.335 |
| Weighted mean |  | 0.634 | **0.567** | 0.616 | 0.621 | 0.610 | 0.605 |
| Unweighted mean |  | 0.575 | **0.499** | 0.519 | 0.523 | 0.511 | 0.507 |

**Table 4.** The kinase inhibition predictive performance comparison

| Client | Test size | Individual MLP model |  |  |  | Union + RFCN | FedAVG + RFCN | FedAMP + RFCN |
| --- | --- | --- | --- | --- | --- | --- | --- | --- |
|  |  | BioMedX | Christmann | PKIS | Tang |  |  |  |
| BioMedX | 170 | 0.648 | 2.533 | 2.634 | 2.189 | **0.490** | 0.766 | 0.611 |
| Christmann | 111 | 0.676 | 0.674 | 0.644 | **0.215** | 0.262 | 1.178 | 0.275 |
| PKIS | 37 | 0.410 | 1.033 | 0.583 | 0.629 | **0.079** | 0.388 | 0.127 |
| Tang | 90 | 0.505 | 0.856 | 1.063 | 0.708 | 0.302 | 0.477 | **0.237** |
| Weighted mean |  | 0.602 | 1.521 | 1.560 | 1.184 | **0.349** | 0.780 | 0.393 |
| Unweighted mean |  | 0.534 | 1.575 | 1.508 | 1.227 | **0.267** | 0.721 | 0.408 |

**Table 5.** The hERG inhibition predictive performance comparison

| Client | Test size | Individual MLP model |  |  |  | Union + MLP | FedAVG + MLP | FedAMP + MLP |
| --- | --- | --- | --- | --- | --- | --- | --- | --- |
|  |  | ChEMBL | NCATS | JHICC | Cai |  |  |  |
| ChEMBL | 748 | 0.842 | 0.639 | 0.074 | 0.839 | **0.892** | 0.871 | 0.882 |
| NCATS | 1652 | 0.453 | 0.697 | 0.014 | 0.546 | **0.752** | 0.686 | 0.632 |
| JHICC | 423 | 0.163 | 0.106 | **0.424** | 0.124 | 0.398 | 0.257 | 0.424 |
| Cai | 786 | 0.837 | 0.628 | 0.025 | 0.810 | **0.885** | 0.873 | 0.882 |
| FDA-Drugs | 177 | **0.516** | 0.457 | 0.021 | 0.504 | 0.442 | 0.395 | 0.428 |
| Keseru | 66 | 0.862 | 0.866 | 0.040 | **0.925** | 0.923 | 0.907 | 0.804 |
| Doddareddy | 1636 | 0.906 | 0.754 | 0.020 | 0.912 | **0.920** | 0.905 | 0.823 |
| Siramshetty | 5804 | 0.966 | 0.658 | 0.024 | 0.890 | **0.976** | 0.958 | 0.868 |
| Zhang | 1565 | 0.821 | 0.698 | 0.022 | 0.864 | **0.881** | 0.861 | 0.754 |
| Sun | 3024 | 0.489 | **0.908** | 0.024 | 0.579 | 0.863 | 0.818 | 0.665 |
| Weighted mean |  | 0.762 | 0.705 | 0.036 | 0.764 | **0.886** | 0.855 | 0.773 |
| Unweighted mean |  | 0.686 | 0.641 | 0.069 | 0.699 | **0.793** | 0.753 | 0.716 |

FedAMP (Federated attentive message passing)^23^, a personalized federated learning aggregation algorithm, is performed here to see how federated learning aggregation algorithm will influence the outcome. FedAMP encourages clients with similar model parameters to have stronger collaboration, so the algorithm adaptively discovers the hidden collaboration relationships between clients and enhancing their collaboration effectiveness by assigning different models to different clients. The combination of FedAMP and RFCN makes it perform best in 4 out of 7 clients, even better than the Union + RFCN baseline. However, in terms of the size-weighted mean and unweighted mean, Union + RFCN model performs the best.

### Case study on kinase inhibition and hERG liability prediction

To demonstrate more use cases, we also simulated federated learning on hERG liability and kinase inhibition data sets. Many kinase inhibitors have the problem of either high toxicity or resistance in tumor^24^. It is of great importance for kinase inhibitors to precisely modulate the wanted kinases as well as avoid the unwanted kinases^25,26^. But usually biotech companies only have the inhibitory activity data of the specific kinase they are developing on. Constructing a predictive model for inhibitory activity across multiple kinases will be helpful for selective inhibitor screening. Federated learning can help them collectively train a more powerful model across multiple kinases. We build a federated model for kinase pIC50 prediction across four data sets from different sources (Supplementary Table 2). As shown in Table 4, the FedAVG + RFCN model perform better than 3 individual models with a large margin but worse than the individual model built on BioMedX dataset. With a better federated aggregation algorithm, FedAMP + RFCN model is better than all of the individual models. However, the best model is the Union + RFCN model trained with mixing data sets in a centralized way in this case.

Drug-induced hERG block is one of the main causes of cardiotoxicity^27^. Assessing hERG liability is required in early drug discovery program. However, various experimental assays can be used to evaluate the hERG liability^28,29^, which induce large biases in the recorded values of hERG liability. Previous studies are focused on merging data from different sources and construct a centralized model to fit the data^30–33^, which can result in biased and overfitted model that may not generalize well. In our study case, a federated hERG classification model was constructed using hERG inhibitory data from different source (Supplementary Table 3). Seen from Table 5, FedAMP + MLP model and FedAMP + MLP model usually outperform modeling on individual data sets but are always inferior to the Union + MLP model.

These two use cases in drug discovery suggest that we can rely on federated learning for better predictive performance without sharing sensitive data, which will largely cut the cost in the help of “knowledge” from each other.

## Discussion

In a bigger federated learning context, the framework we set up only focuses on simulating participants who have the same feature space (molecular ECFP fingerprints) as input, which is attributed to horizontal federated learning (Supplementary Figure 1A)^34^. There is also vertical federated learning scheme can cope with participants having different feature types as input (Supplementary Figure 1B). Moreover, a combination of horizontal and vertical federated learning can effectively handle the participants who share some feature types and samples but also have their own proprietary feature types and samples, which is referred to as federated transfer learning^34^. Federated transfer learning will further expand the feature space and sample size we have by taking both the union of feature space and sample space of multiple participants. For example, to predict the clinical outcome of drug candidates, we need integrate data with shared and proprietary features from multiple parties, including related pharmaceutical companies, hospitals and patients, federated transfer learning may generate large adding value for each party.

Federated learning may still get some security concerns and malicious or non-malicious failures, but it has attracted substantial attention and has been improving and evolving quickly. This paradigm opens up the possibility to integrate confidential datasets through secure distributed training, which have previously been considered impractical but absolutely attractive to drug discovery. Given the predictive model in drug discovery often work in very confined domain, the opportunity to leverage larger and diverse data silos from multiple institutions will improve generalizability of the predictive models in drug discovery.

## Conclusion

In this work, we set up a cross-silo federated learning framework for drug discovery based on FATE^19^, and constructed baseline models using MLP and RFCN architectures. We collected 7 drug solubility datasets and simulated the whole process including federated training, model selection, rollout and customization. Federated training can perform better than individual training on each dataset and, more surprisingly, better than centralized training on the pooled highly biased datasets. Visualizing the weight distributions of parameters in the neural network, we find federated training learned a simpler model with more zero weights than conventional centralized training, which means federated learning intrinsically has a regularization effect and may contribute to better generalization performances in highly biased data. Beyond that, it is feasible for federated learning to customize the global model locally (don’t need to upload the model updates) if new users having plenty of labeled data. We demonstrated that users can get some benefits from customizing the global model than using the global model directly. Federated learning represents a new machine learning paradigm, the feature of privacy-preserving will encourage more and more institutions to fully utilize their data and expose more and more data to the latest machine learning models, thus solving the “small data” dilemma in drug discovery. Federated learning setting also makes it feasible to customize models for different users/clients, and hence alleviate the problem of data bias and achieve better predictive performance and form wiser instructions in real application scenario.

## Methods

### Data curation and partitioning

The 7 aqueous solubility datasets are collected from 7 different sources, which are preprocessed and curated by Sorkun et al. in AqSolDB^35^. Dataset F1 was extracted from eChemPortal, an open-source chemical property database developed by the OECD (Organisation for Economic Co-operation and Development)^36^. Both Dataset F2 and Dataset F4 are obtained EPI Suite Data website, which were generated by Water Solubility Fragment program^37^ and WSKOWWIN program^38^, respectively. Dataset F3 was taken from the work of Raevsky et al.^39^. Dataset C1 was collected from the work of Huuskonen et al.^40^. Dataset C2 was collected from the work of Wang et al.^41^. Dataset C3 was collected from the work of Delaney et al.^42^.

In our simulation, the owners of datasets F1-4 are supposed to be the collaborated parties who want to participate in federated training, and the owners of datasets C1-3 are users who want to use the federated trained model. To prevent overfitting, 1/10 molecules of datasets F1-4 were held out as validation set. Another 1/10 molecules were held out as test sets, for comparing with different models. Because user C1-3 have their own data, it is feasible for them to customize their own model by fine tuning the federated trained model. We set the proportion of train, validation and test sets of C1-3 by 3:1:1.

Kinase inhibition datasets were curated by Merget and Fulle et al. and taken from https://github.com/Team-SKI/Publications^43^, which contains four datasets—the Tang set^44^ (a collection of the kinase profiling data sets of Metz^45^, Davis^46^ and Anastassiadis^47^), PKIS^48–50^, Christmann2016^51^ and a curated ChEMBL kinase inhibitor panel by Merget^52^. All these four datasets are used for federated training.

hERG liability datasets are collected from different sources that include Cai et. al.^53^, Zhang et al.^54^, Siramshetty et al.^55^, Keseru et al.^30^, Doddareddy et al.^56^, Sun et al.^57^, Pubchem NCATS^58^, Pubchem JHICC^59^, ChEMBL^60^ and FDA-Drugs^61^. Datasets from Cai et al., Pubchem NCATS, Pubchem JHICC and ChEMBL are simulated to be the clients who take part in federated training, the rest of those datasets are only simulated as test sets.

### Federated Averaging and Secure Aggregation

In our setting, all clients have the same features (molecular fingerprints) as input for prediction task, so all the clients are deployed with the same neural networks architecture and could be trained with Federated Averaging. To ensure the security of data and model, the model updates should also not be uploaded in plaintext. Therefore, a Secure Aggregation protocol are implemented together with Federated Averaging. Both the Federated Averaging and Secure Aggregation protocol are proposed by Google’s team in separate works^7,62^.

As descripted in pseudo-code of Algorithm 1, when the training start, the coordinator initializes the model parameters *w*_0_, which will be broadcast to each client. In each round of federated training, the client downloads the current shared global model *w*_*t*_ from the coordinator server, and trains its model locally on its own data with SGD optimizer. At every E (a hyperparameter) epochs of local training, all the clients will compute the updates and encrypted them under Secure Aggregation protocol. As per the protocol, the local model updates of each client would be added a unique random mask that are carefully generated and relevant to all the other participants, so as to make sure all the random masks adds up to 0 and thus be cancelled out when the coordinator aggregates the local updates uploaded by all clients. Since the random masks are cancelled out, the coordinator gets the true averaged model updates and uses them to update the federated model parameters, obtaining the current shared model. The shared global model will be broadcast to all clients, starting a new round of training. Similar to conventional neural networks, the training process will stop when the federated model converges or the training process reaches a predefined max-round threshold. Note that not as simple as descripted in Algorithm 1, Secure Aggregation Protocol is much more complicated with a four-round interaction between the coordinator and clients, which make protocol robust to dropouts and delays of the clients. The Federated Averaging and Secure Aggregation Protocol are implemented on FATE^19^.

### Federated attentive message passing

Most of the existing federated learning practices are not able to achieve good performances because a single global model is used for all clients. Personalized federated learning allows us to train a personalized model without leaking the private data. FedAMP (Federated attentive message passing)^23^, a personalized federated learning aggregation algorithm, has not implemented on FATE and we simulated the process.

#### Algorithm 1 Federated averaging with secure aggregation

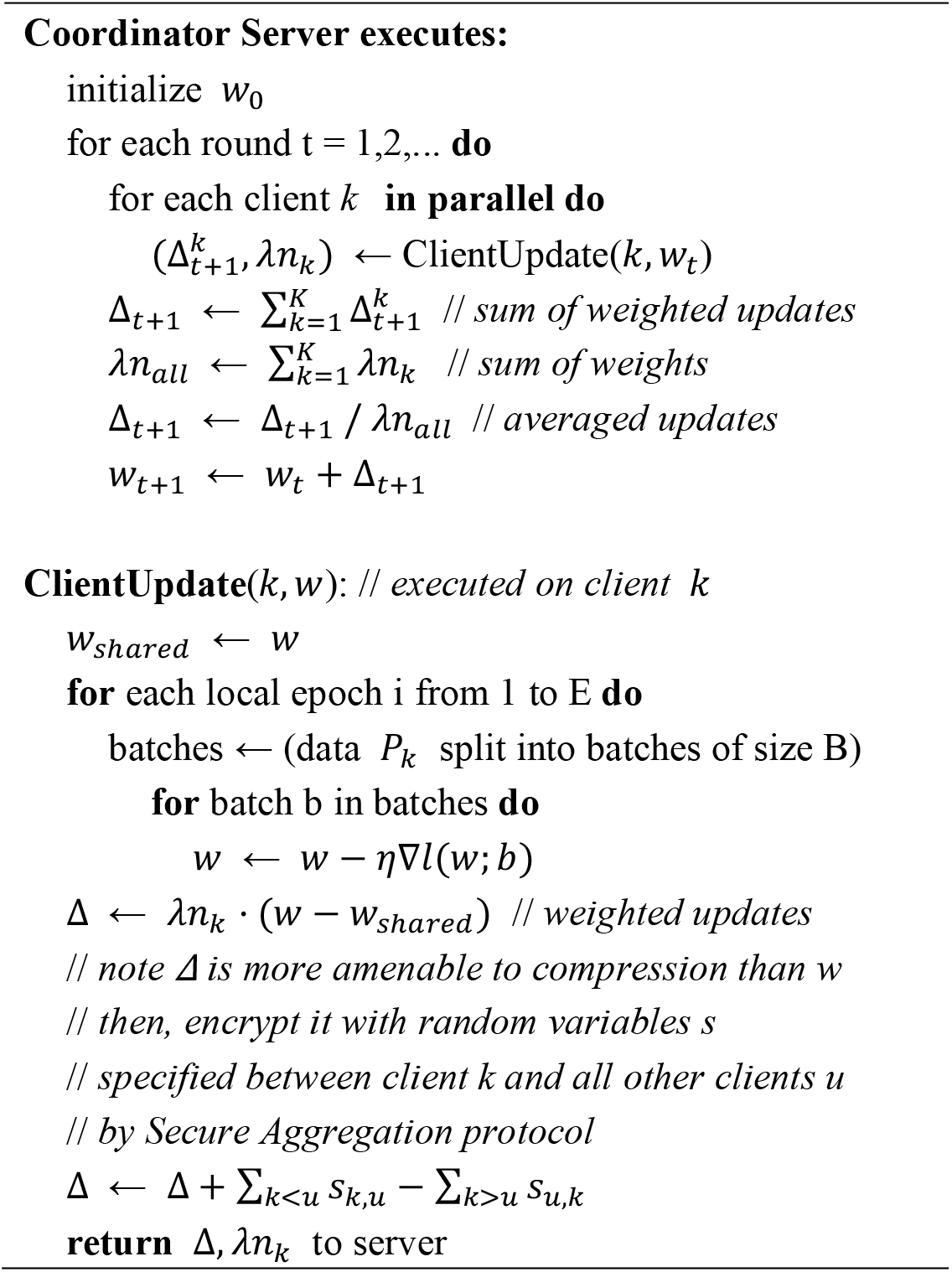

### Neural network architecture and training

As illustrated in Figure 2B, for the aqueous solubility datasets, MLP model takes the 2048 bit ECFP molecular fingerprints^63^ as input and goes through 6 fully connected layers activated with the relu function, between which there are 4 batch normalization layers and two dropout layers with dropout rates of 0.2, 0.3, respectively^20^. The RFCN is composed of a fully connected layer with 1536 neurons followed by a ResNet of two 2-layer blocks and one single-layer block (Supplementary Figure 2B). The network architecture is shared by all the clients. All the models are trained with backpropagation and SGD optimizer, where the learning rate starts with 0.01 and follows a 0.68 decay rate^64^. For the kinase inhibitory activity datasets, the RFCN consists of two fully connected layers and a DenseNet of two 2-layer dense blocks. For the hERG liability datasets, the MLP is composed of a 6-layer multilayer perceptron with neuron numbers of 4096, 1024, 256, 64, 6 and 2 respectively.

### Model selection and personalization

The model is selected by looking at the averaged MAE of the held out validation set in the training participants, if it doesn’t improve in 30 rounds, the training would stop. When the best shared global federated model is selected, it can be used for prediction directly. However, new users who did not participate in the federated training may personalize their own model by getting the licenses and instrumenting the federated training program as done by the participants of federated training. Then, they can fine tune the federated model locally on their own labeled data without uploading the model updates.

## AUTHOR INFORMATION

### Notes

The authors declare no competing financial interest.

## ACKNOWLEDGMENT

We gratefully acknowledge financial support from the National Natural Science Foundation of China (81773634 to M.Z), National Science & Technology Major Project “Key New Drug Creation and Manufacturing Program”, China (Number: 2018ZX09711002 to H.J.), and “Personalized Medicines—Molecular Signature-based Drug Discovery and Development”, Strategic Priority Research Program of the Chinese Academy of Sciences (XDA12050201 and XDA12020368).

## Code and data availability

The data and federated model built on FATE are available at https://github.com/chengziqiang/FL_DrugDiscoverry

## Reference

1. Smalley, E. AI-powered drug discovery captures pharma interest. Nat. Biotechnol. 35, 604–605 (2017).

2. Schneider, P. et al. Rethinking drug design in the artificial intelligence era. Nat. Rev. Drug Discov. (2019) doi: 10.1038/s41573-019-0050-3.

3. Chen, B. et al. Harnessing big ‘omics’ data and AI for drug discovery in hepatocellular carcinoma. Nat. Rev. Gastroenterol. Hepatol. (2020) doi: 10.1038/s41575-019-0240-9.

4. Hunter, A. J., Lee, W. H. & Bountra, C. Open innovation in neuroscience research and drug discovery. Brain Neurosci. Adv. 2, 2398212818799270 (2018).

5. Zhou, Y. Drug development and medical writing in the digital world. Med. Writ. 28, 18–21 (2019).

6. Riley, P. Three pitfalls to avoid in machine learning. Nature 572, 27–29 (2019).

7. McMahan, B., Moore, E., Ramage, D., Hampson, S. & y Arcas, B. A. Communication-Efficient Learning of Deep Networks from Decentralized Data. in Artificial Intelligence and Statistics 1273–1282 (2017).

8. Yang, T. et al. Applied Federated Learning: Improving Google Keyboard Query Suggestions. ArXiv E-Prints 1812.02903 (2018).

9. Bonawitz, K. et al. Towards Federated Learning at Scale: System Design. ArXiv E-Prints 1902.01046 (2019).

10. Kairouz, P. et al. Advances and open problems in federated learning. ArXiv Prepr. 191204977 (2019).

11. Wang, K. et al. Federated Evaluation of On-device Personalization. ArXiv E-Prints 1910.10252 (2019).

12. Jiang, Y., Konečný, J., Rush, K. & Kannan, S. Improving Federated Learning Personalization via Model Agnostic Meta Learning. ArXiv E-Prints 1909.12488 (2019).

13. WeBank. WeBank and Swiss signed cooperation MOU. https://finance.yahoo.com/news/webank-swiss-signed-cooperation-mou-112300218.html (2019).

14. Li, W. et al. Privacy-Preserving Federated Brain Tumour Segmentation. in Machine Learning in Medical Imaging (eds. Suk, H.-I., Liu, M., Yan, P. & Lian, C.) 133–141 (Springer International Publishing,2019).

15. FeatureCloud. FeatureCloud: Our vision. https://featurecloud.eu/about/our-vision/ (2019).

16. Musketeer. Musketeer: About. http://musketeer.eu/project/ (2019).

17. ai.intel. Federated learning for medical imaging. https://www.intel.ai/federated-learning-for-medical-imaging/ (2019).

18. Cordis, E. Machine learning ledger orchestration for drug discovery. https://cordis.europa.eu/project/rcn/223634/factsheet/en?WT.mc_id=RSS-Feed&WT.rss_f=project&WT.rss_a=223634&WT.rss_ev=a (2019).

19. WeBank. FATE (Federated AI Technology Enabler). https://github.com/FederatedAI/FATE.

20. Srivastava, N., Hinton, G., Krizhevsky, A., Sutskever, I. & Salakhutdinov, R. Dropout: A simple way to prevent neural networks from overfitting. J Mach Learn Res 15, 1929 (2014).

21. Smirnov, E. A., Timoshenko, D. M. & Andrianov, S. N. Comparison of Regularization Methods for ImageNet Classification with Deep Convolutional Neural Networks. 2nd AASRI Conf. Comput. Intell. Bioinforma. 6, 89–94 (2014).

22. Liu, D. et al. AutoGenome: An AutoML Tool for Genomic Research. bioRxiv (2019) doi: 10.1101/842526.

23. Huang, Y. et al. Personalized Federated Learning: An Attentive Collaboration Approach. (2020).

24. Zhang, W., Roederer, M. W., Chen, W.-Q., Fan, L. & Zhou, H.-H. Pharmacogenetics of drugs withdrawn from the market. PHARMACOGENOMICS 13, 223–231 (2012).

25. Ai, X., Sun, Y., Wang, H. & Lu, S. A Systematic Profile of Clinical Inhibitors Responsive to EGFR Somatic Amino Acid Mutations in Lung Cancer: Implication for the Molecular Mechanism of Drug Resistance and Sensitivity. Amino Acids 46, 1635 (2014).

26. Daub, H., Specht, K. & Ullrich, A. Strategies to overcome resistance to targeted protein kinase inhibitors. Nat. Rev. Drug Discov. 3, 1001–1010 (2004).

27. Anwar-Mohamed, A. et al. A human ether-a-go-go-related (hERG) ion channel atomistic model generated by long supercomputer molecular dynamics simulations and its use in predicting drug cardiotoxicity. Toxicol. Lett. 230, 382–392 (2014).

28. Aronov, A. & Goldman, B. A model for identifying HERG K+ channel blockers. Bioorg. Med. Chem. 12, 2307–2315 (2004).

29. Beaugrand, M., Arnold, A. A., Bourgault, S., Williamson, P. T. F. & Marcotte, I.Comparative study of the structure and interaction of the pore helices of the hERG and Kv1.5 potassium channels in model membranes. Eur. Biophys. J. Biophys. Lett. 46, 549–559 (2017).

30. Keseru, G. M. Prediction of hERG potassium channel affinity by traditional and hologram QSAR methods. Bioorg. Med. Chem. Lett. 13, 2773–2775 (2003).

31. Aronov, A. M. Predictive in silico modeling for hERG channel blockers. Drug Discov. Today 10, 149 (2005).

32. Benson, A. P., Al-Owais, M. & Holden, A. V. Quantitative prediction of the arrhythmogenic effects of de novo hERG mutations in computational models of human ventricular tissues. Eur. Biophys. J. Biophys. Lett. 40, 627–639 (2011).

33. Braga, R. C. et al. Pred-hERG: A Novel web-Accessible Computational Tool for Predicting Cardiac Toxicity. Mol. Inform. 34, 698–701 (2015).

34. Yang, Q., Liu, Y., Chen, T. & Tong, Y. Federated Machine Learning: Concept and Applications. ArXiv E-Prints 1902.04885 (2019).

35. Sorkun, M. C., Khetan, A. & Er, S. AqSolDB, a curated reference set of aqueous solubility and 2D descriptors for a diverse set of compounds. Sci. Data 6, 143 (2019).

36. OECD. eChemPortal - The Global Portal to Information on Chemical Substances. https://www.echemportal.org/echemportal/propertysearch/addblock_input.action.

37. US EPA. EPI Suite Data. WATERNT (Water Solubility Fragment) Program Methodology & Validation Documents,. http://esc.syrres.com/interkow/Download/WaterFragmentDataFiles.zip.

38. US EPA. EPI Suite Data. WSKOWWIN Program Methodology & Validation Documents. http://esc.syrres.com/interkow/Download/WSKOWWIN_Datasets.zip.

39. Raevsky, O. A., Grigor’ev, V. Yu., Polianczyk, D. E., Raevskaja, O. E. & Dearden, J. C. Calculation of Aqueous Solubility of Crystalline Un-Ionized Organic Chemicals and Drugs Based on Structural Similarity and Physicochemical Descriptors. J. Chem. Inf. Model. 54, 683–691 (2014).

40. Huuskonen, J. Estimation of aqueous solubility for a diverse set of organic compounds based on molecular topology. J Chem Inf Comput Sci 40, 773 (2000).

41. Wang, J., Hou, T. & Xu, X. Aqueous Solubility Prediction Based on Weighted Atom Type Counts and Solvent Accessible Surface Areas. J. Chem. Inf. Model. 49, 571–581 (2009).

42. Delaney, J. S. ESOL: estimating aqueous solubility directly from molecular structure. J Chem Inf Comput Sci 44, 1000 (2004).

43. Merget, B., Turk, S., Eid, S., Rippmann, F. & Fulle, S. Profiling Prediction of Kinase Inhibitors: Toward the Virtual Assay. J. Med. Chem. 60, 474–485 (2017).

44. Tang, J. et al. Making sense of large-scale kinase inhibitor bioactivity data sets: a comparative and integrative analysis. J Chem Inf Model 54, 735 (2014).

45. Metz, J. T. et al. Navigating the kinome. Nat Chem Biol 7, 200 (2011).

46. Davis, M. I. et al. Comprehensive analysis of kinase inhibitor selectivity. Nat Biotechnol 29, 1046 (2011).

47. Anastassiadis, T., Deacon, S. W., Devarajan, K., Ma, H. & Peterson, J. R. Comprehensive assay of kinase catalytic activity reveals features of kinase inhibitor selectivity. Nat Biotechnol 29, 1039 (2011).

48. Dranchak, P. et al. Profile of the GSK published protein kinase inhibitor set across ATP-dependent and-independent luciferases: implications for reporter-gene assays. PLoS One 8, e57888 (2013).

49. Knapp, S. et al. A public-private partnership to unlock the untargeted kinome. Nat Chem Biol 9, 3 (2013).

50. Elkins, J. M. et al. Comprehensive characterization of the Published Kinase Inhibitor Set. Nat Biotechnol 34, 95 (2015).

51. Christmann-Franck, S. et al. Unprecedently Large-Scale Kinase Inhibitor Set Enabling the Accurate Prediction of Compound-Kinase Activities: A Way toward Selective Promiscuity by Design? J. Chem. Inf. Model. 56, 1654–1675 (2016).

52. Volkamer, A. et al. Pocketome of human kinases: prioritizing the ATP binding sites of (yet) untapped protein kinases for drug discovery. J Chem Inf Model 55, 538 (2015).

53. Cai, C. et al. Deep Learning-Based Prediction of Drug-Induced Cardiotoxicity. J. Chem. Inf. Model. 59, 1073–1084 (2019).

54. Zhang, S. T., Zhou, Z. F., Gong, Q. M., Makielski, J. C. & January, C. T. Mechanism of block and identification of the verapamil binding domain to HERG potassium channels. Circ. Res. 84, 989–998 (1999).

55. Siramshetty, V. B. et al. Critical Assessment of Artificial Intelligence Methods for Prediction of hERG Channel Inhibition in the ‘Big Data’Era. (2020).

56. Doddareddy, M. R., Klaasse, E. C., Shagufta, Ijzerman A. P. & Bender, A. Prospective Validation of a Comprehensive In silico hERG Model and its Applications to Commercial Compound and Drug Databases. CHEMMEDCHEM 5, 716–729 (2010).

57. Sun, X. et al. Characterization and structure-activity relationship of natural flavonoids as hERG K+ channel modulators. Int. Immunopharmacol. 45, 187–193 (2017).

58. Pubchem NCATS. https://pubchem.ncbi.nlm.nih.gov/bioassay/588834 (2019).

59. Pubchem JHICC. https://pubchem.ncbi.nlm.nih.gov/bioassay/2321 (2019).

60. Bento, A. P. et al. The ChEMBL bioactivity database: an update. Nucleic Acids Res 42, D1083 (2014).

61. Drugs@FDA: FDA-Approved Drugs. https://www.accessdata.fda.gov/scripts/cder/daf/index.cfm (2019).

62. Bonawitz, K. et al. Practical Secure Aggregation for Privacy-Preserving Machine Learning.in Proceedings of the 2017 ACM SIGSAC Conference on Computer and Communications Security 1175–1191 (Association for Computing Machinery, 2017). doi: 10.1145/3133956.3133982.

63. Rogers, D. & Hahn, M. Extended-connectivity fingerprints. J Chem Inf Model 50, 742 (2010).

64. Haddadpour, F., Mahdi Kamani, M., Mahdavi, M. & Cadambe, V. R. Local SGD with Periodic Averaging: Tighter Analysis and Adaptive Synchronization. ArXiv E-Prints 1910.13598 (2019).

